# System Identification Using Compressed Sensing Reveals Signaling-Decoding System by Gene Expression

**DOI:** 10.1101/129296

**Authors:** Takaho Tsuchiya, Masashi Fujii, Naoki Matsuda, Katsuyuki Kunida, Shinsuke Uda, Hiroyuki Kubota, Katsumi Konishi, Shinya Kuroda

**Author notes:** Lead Contact: Shinya Kuroda, Hongo 7-3-1, Bunkyo-ku, Tokyo 113-0033, Japan.

## Abstract

Cells decode information of signaling activation at a scale of tens of minutes by downstream gene expression with a scale of hours to days, leading to cell fate decisions such as cell differentiation. However, no system identification method with such different time scales exists. Here we used compressed sensing technology and developed a system identification method using data of different time scales by recovering signals of missing time points. We measured phosphorylation of ERK and CREB, immediate early gene expression products, and mRNAs of decoder genes for neurite elongation in PC12 cell differentiation and performed system identification, revealing the input–output relationships between signaling and gene expression with sensitivity such as graded or switch-like response and with time delay and gain, representing signal transfer efficiency. We predicted and validated the identified system using pharmacological perturbation. Thus, we provide a versatile method for system identification using data with different time scales.

**Highlights:**

- We developed a system identification method using compressed sensing.
- This method allowed us to find a pathway using data of different time scales.
- We identified a selective signaling-decoding system by gene expression.
- We validated the identified system by pharmacological perturbation.

**eTOC Blurb:** We describe a system identification method of molecular networks with different time-scale data using a signal recovery technique in compressed sensing.

## INTRODUCTION

In intracellular signaling systems, information of an extracellular stimulus is encoded into combinations of distinct temporal patterns of phosphorylation of intracellular signaling molecules that are selectively decoded by downstream gene expression, leading to cell fate decisions such as cell differentiation, proliferation, and death (Behar and Hoffmann, 2010; Purvis and Lahav, 2013). For instance, in rat adrenal pheochromocytoma PC12 cells, nerve growth factor (NGF) induces cell differentiation mainly through sustained phosphorylation of ERK (Gotoh et al., 1990; Marshall, 1995; Qiu and Green, 1992; Traverse et al., 1992), whereas pituitary adenylate cyclase-activating polypeptide (PACAP) induces cell differentiation mainly through cAMP-dependent CREB phosphorylation (Akimoto et al., 2013; Gerdin and Eiden, 2007; Saito et al., 2013; Vaudry et al., 2002; Watanabe et al., 2012). We showed that cell differentiation in PC12 cells can be divided into two processes: a latent processes (0–12 h after the stimulation) in preparation for neurite extension and a subsequent neurite extension process (12–24 h) (Chung et al., 2010). We identified the three genes essential for cell differentiation, *Metrnl*, *Dclk1*, and *Serpinb1a*, which are induced during the latent process and required for subsequent neurite extension, and named LP (latent process) genes (Watanabe et al., 2012). Although NGF and PACAP selectively induce the different combinations and temporal patterns of signaling molecules, both growth factors commonly induce the LP genes (Watanabe et al., 2012). The expression levels of LP genes, but not the phosphorylation level of ERK, correlate with neurite length regardless of growth factors (Watanabe et al., 2012), indicating that the LP genes are the decoders of neurite length. Thus, how the distinct patterns of signaling molecules are decoded by LP gene expression is critical for understanding the unknown mechanism underlying cell differentiation in PC12 cells. Decoding the combinations and temporal patterns of signaling molecules by downstream gene expression is a general mechanism underlying various cellular functions (Behar and Hoffmann, 2010; Purvis and Lahav, 2013; Sumit et al., 2017).

Mathematical modeling is useful for the analysis of decoding mechanisms (Janes and Lauffenburger, 2013). If the signaling pathways are well characterized, kinetic modeling based on biochemical reactions reported in the literature is often used (Janes and Lauffenburger, 2006; Kholodenko et al., 2012; Price and Shmulevich, 2007). For example, growth factor–dependent ERK activation in PC12 cells has been modelled by the kinetic model based on prior knowledge of pathway information (Brightman and Fell, 2000; Filippi et al., 2016; Nakakuki et al., 2010; Ryu et al., 2015; Santos et al., 2007; Sasagawa et al., 2005). In general, however, decoding by downstream genes involves more complex processes such as transcription and translation and information on the precise pathway is not available.

To identify decoding mechanisms by gene expression, the system identification method (also referred to as data-driven modeling) was developed for identifying quantitative input–output relationships from time series data without detailed knowledge of signaling pathways (Janes and Lauffenburger, 2006; Janes and Yaffe, 2006; Kholodenko et al., 2012; Ljung, 2010; Price and Shmulevich, 2007; Zechner et al., 2016). We previously developed a system identification method based on time series data of signaling molecules and gene expression, denoted as the nonlinear autoregressive exogenous (NARX) model, and applied it to the signaling-dependent immediate early gene (IEG) expression during cell differentiation in PC12 cells (Saito et al., 2013). The NARX model involves the determination of lag-order numbers and use of the Hill equation and the linear autoregressive exogenous (ARX) model (Saito et al., 2013).

Determination of lag-order numbers infers the selection of input molecules (*Input*) for an output molecule (*Output)*, which is referred to as the *Input*-*Output* (*I*-*O*). The Hill equation characterizes sensitivity with a nonlinear dose-response curve (Hill, 1910). The linear ARX model characterizes temporal changes with time constant and gain, the latter of which is an *I*-*O* amplitude ratio, and indicates signal transfer efficiency (Ljung, 1998). The NARX model requires equally spaced dense time series data. If the time scale between upstream and downstream are similar, such as signaling molecules (scale of tens of minutes) and IEG expression (a few hours) in PC12 cells, it is not difficult to acquire a sufficient number of equally spaced dense time series data (Saito et al., 2013). However, if the time scale of upstream and downstream molecules is largely different, such as signaling molecules (tens of minutes) and LP gene expression (a day) (Doupé and Perrimon, 2014), it is technically difficult to obtain sufficient equally spaced dense time series data because of experimental and budget limitations.

Measuring gene expression often requires a longer time scale than measuring protein phosphorylation. Obtaining equally spaced dense time series data with a longer time scale is labor and cost intensive, because, unlike live-cell imaging experiments, snapshot experiments such as western blotting, RT-PCR, and quantitative image cytometry (QIC) (Ozaki et al., 2010) require the same number of experiments as the number of time points. In addition, experimental noise and variation increases as the number of experiments increases because differences in experimental conditions such as plates, gels, reagents, and cell culture conditions increase as well. Therefore, in reality, for a longer time scale experiment, unequally spaced sparse time series data rather than equally spaced dense time series data are desired. For example, under conditions in which stimulation by cell growth factors triggers rapid and transient phosphorylation and slow and sustained gene expression, time series data should be obtained with dense time points during the transient phase and eventually with sparse time points. The timing and dynamic characteristics of temporal changes may differ between upstream and downstream molecules, such that time points and intervals for measuring upstream and downstream molecules may be different. Thus, a system identification method using unequally spaced sparse time series data with different time scale needs to be developed.

To solve this problem, here we used the signal recovery technique based on a low-rank approach proposed in the field of compressed sensing to generate a sufficient number of time points for equally spaced dense time series data from unequally spaced sparse time series data with different time points and intervals. We applied this nonlinear system identification method to the signaling-dependent gene expression underlying cell differentiation in PC12 cells and identified the signaling-decoding system by gene expression.

Unequally spaced sparse time series data can be regarded as equally spaced dense time series data with missing time points, and therefore we can generate equally spaced dense time series data by applying a signal recovery technique, which has been studied in the field of compressed sensing (Candès and Wakin, 2008; Donoho, 2006). Compressed sensing is a signal processing method for efficient data acquisition by recovering missing signals/images from a small number of randomly sampled signals including unequally spaced sparse data based on sparseness of a vector (Candès et al., 2008) or low rankness of a matrix (Fazel, 2002). Both the sparse approach and the low-rank approach have been applied to various fields, such as sampling and reconstructing magnetic resonance images (Lustig et al., 2008; Ongie and Jacob, 2016), super-resolution imaging (Candès and Fernandez-Granda, 2014; Yang et al., 2010), image inpainting (Takahashi et al., 2012; Takahashi et al., 2016), and collaborative filtering (Candès and Recht, 2009). In this study, we applied a matrix rank minimization algorithm (Konishi et al., 2014) to recover missing time points from unequally spaced time series data, and we generated equally spaced time series data with the same time points from signaling and gene expression data with different time scales. We previously developed a system idenfitication method from equally spaced dense time series data of signaling and gene expression using the NARX model (Saito, 2013). We developed a new system identification method from unequally spaced sparse time series data with different time scales by integrating this signal recovery method using the matrix rank minimization algorithm (Konishi et al., 2014) and the NARX model (Saito et al., 2013). We applied the method to system identification of signaling-dependent gene expression in cell differentiationin PC12 cells, revealing a selective signaling-decoding mechansim by gene expression.

## RESULTS

### Signal Recovery Using Compressed Sensing from Unequally Spaced Data

In this study, we regarded unequally spaced sparse time series data as equally spaced dense time series data with missing time points, and equally spaced time series data were generated by restoring missing time points using a low-rank approach (Konishi et al., 2014). In the low-rank approach for image recovery, we assumed that the value of each pixel is represented by a linear combination of its neighbor pixels, which is mathematically represented by an autoregressive (AR) model. Then a Hankel-like matrix composed of pixel values has a low rank because each column is represented by the linear combination of the other columns (Figures 1 and S1A). This means that the Hankel-like matrix is a low-rank matrix whose rank is determined by the system order. Missing data can be recovered by estimating missing elements of the matrix so that the rank of this matrix [*Y*] is *r*. When system order *r* is unknown, based on the idea that the system can be described with as few parameters as possible, missing elements of this Hankel-like matrix are recovered so as to minimize the rank of the matrix [*Y*]. Based on the low rankness of the Hankel matrix, the signal recovery problem of the missing pixles can be formulated as a matrix rank minimization problem, and we can restore an image by solving this problem (Takahashi et al., 2012; Takahashi et al., 2016) (Figure 1).

**Figure 1.**
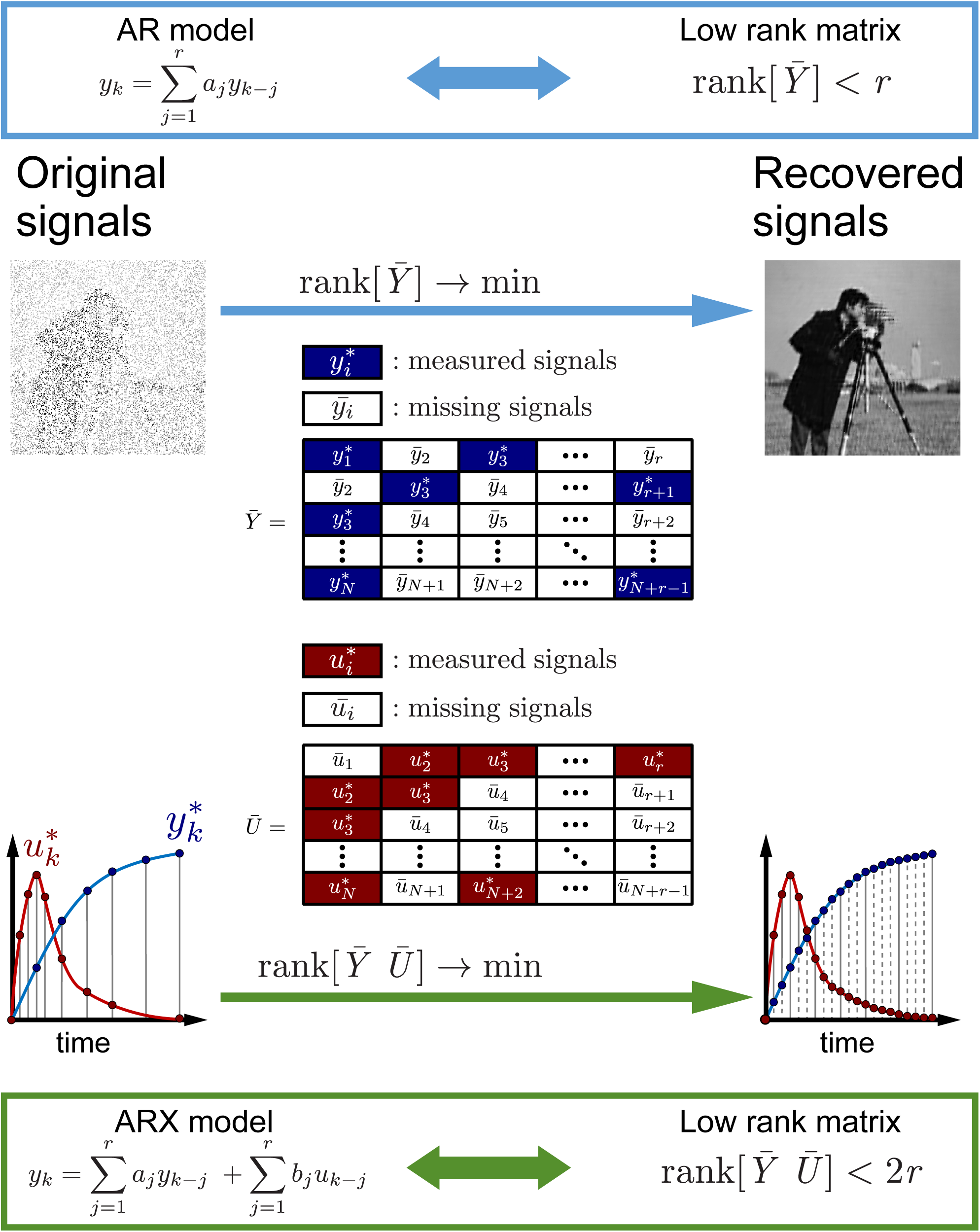
Signal recovery based on compressed sensing technology from unequally spaced data. (Top) An example of recovered equally spaced image data from unequally spaced image data by the signal recovery technique using rank minimization of the Hankel-like matrix ***Y*** composed of signals in pixels. We assume that the value of each pixel is represented by a linear combination of those of its neighboring pixels, which is mathematically represented by an autoregressive (AR) model. (Bottom) An example of recovered equally spaced time series data from unequally spaced time series data by the signal recovery technique using rank minimization of the Hankel-like matrices ***Y*** and ***U***, composed of time series data of input molecules (*Inputs*) and output molecules (*Outputs*), respectively. We assume that the value at a certain time is represented not only by the linear combination of values of the output molecule at past points but also by the linear combination of the values of the input molecule at past points, which is mathematically represented by an autoregressive exogenous (ARX) model. The recovered time series input–output data have the equally spaced time series data with the same time points even if the missing time points of input and output are different.

We performed system identification from unequally spaced time series data of input molecules (*Inputs*) and output molecules (*Outputs*). Although an AR model is used for image recovery, we used an ARX model where the value at a time point is represented by a linear combination of two kinds of signals, *Inputs* and *Outputs*. Therefore, we modified the rank-minimization-based signal recovery method of the AR model to the ARX model and performed system identification (Figures 1 and S1B). Several methods for system identification using a linear ARX model with signal recovery of missing points of input and output based on matrix rank minimization have been proposed (Liu et al., 2013; Markovsky, 2013). They can recover missing time series input–output data even when missing time points of input are not equal to those of output.

However, we cannot directly apply the method because we used the NARX model rather than the ARX model due to the nonlinearity of signaling-dependent gene expression (Kudo et al., 2016; Saito et al., 2013). Therefore, by combining the nonlinear ARX system identification method (Saito et al., 2013) and the signal recovery method based on the matrix rank minimization problem (Konishi et al., 2014), we derived the signal recovery algorithm applicable to the nonlinear ARX system and performed system identification using recovered equally spaced time series input–output data (see “NARX Model and Data Representation” and “Extension ARX System Identification from Unequally Spaced Time Series Data to the NARX System” sections in the STAR METHODS).

### System Identification by Integrating Signal Recovery and the NARX Model

In the NARX model used in our previous work, time series data of *Inputs* are nonlinearly transformed using the Hill equation, which are then used as inputs for the ARX model (Saito et al., 2013) (Figure 2A). The Hill equation, which is nonlinear transformation function *f*(*x*) widely used in biochemistry (Hill, 1910), can represent sensitivity with a graded or switch-like response by the values of *n* and *K* (Figure 2A). The ARX model in the NARX model can represent how the *Output* efficiently responds to the temporal change of the nonlinearly transformed *Inputs* by the time constant and gain (Figure 2A). Thus, from the estimated parameters of the Hill equation and ARX model, the sensitivity with graded or switch-like response and the time constant and gain are obtained, respectively. In this study, the parameters of this NARX model were estimated using a signal recovery scheme based on a low-rank approach (Konishi et al., 2014), as follows (Figure 2B, see details in “Procedure for System Identification by Integrating Signal Recovery and the NARX Model” section in the STAR METHODS).

**Figure 2.**
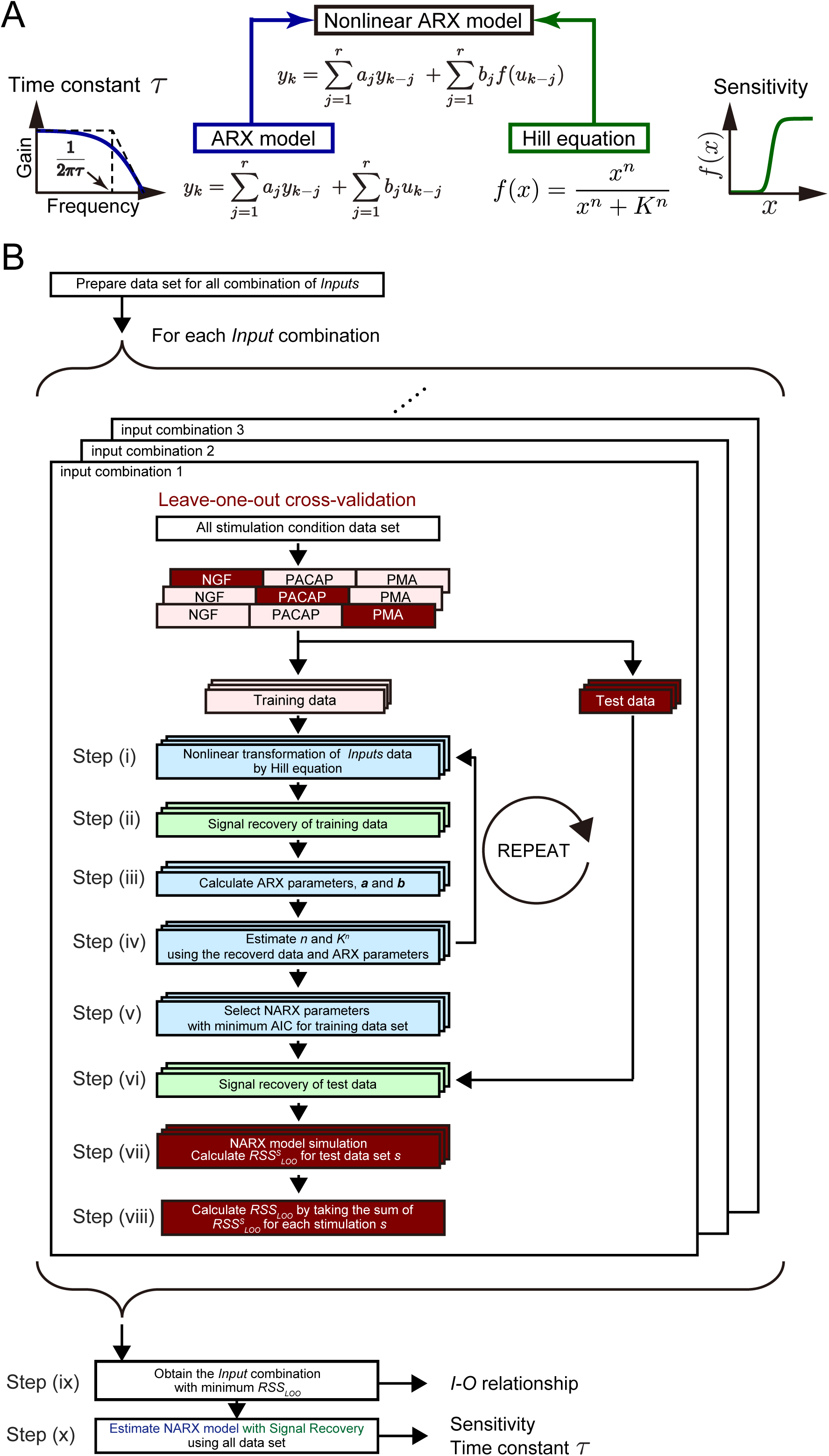
System identification by integrating signal recovery and the NARX model. (A) The nonlinear ARX (NARX) model consists of static nonlinear conversion of input signal by the Hill equation, followed by time delay by ARX model (Saito et al., 2013). The former gives the sensitivity with a graded or switch-like response and the latter gives the time constant. (B) Algorithm flowchart for system identification by integrating signal recovery and the NARX model. See details in “Procedure for System Identification by Integrating Signal Recovery and the NARX Model” section in the STAR METHODS.

To estimate the *I*-*O* relationship, we selected a combination of *Inputs* for each *Output* and prepared a data set of all combinations of *Inputs* for each *Output*. The each data set was divided into test dataset for one stimulation condition and training data set for the rest of two stimulation conditions, leave-one-out (LOO) cross-validation was performed. We estimated the parameters of the NARX model (the NARX parameters) for the training data set in each *Input–Output* combination using the following method. First, the initial values of *n* and *K* are given by *n* = 1 and a random number, respectively, and nonlinear transformation of input unequally spaced time series data by using the Hill equation was performed (Figure 2B, step i). A Hankel-like matrix was constructed from unequally spaced time series *Output* data and from unequally spaced time series *Inputs* nonlinearly transformed by the Hill equation. Next, signal recovery was performed with an iterative partial matrix shrinkage (IPMS) algorithm to minimize the rank of the Hankel-like matrix composed of *Output* and *Inputs* transformed by the Hill equation (Konishi et al., 2014) (Figure 2B, step ii). The rank of the recovered Hankel-like matrix corresponds to the lag order; from the recovered Hankel-like matrix, the parameters of the ARX model ***a*** and ***b*** were uniquely obtained (Figure 2B, step iii; Figure S1B). Further estimation of *n* and *K* was performed by using recovered data, ARX parameters obtained until step iii, and other combination of *n* and *K* given random numbers (Figure 2B, step iv). By using the inverse function of the Hill equation, we recovered the missing time points data of input before transformation by the Hill equation. For the other 200 combinations of *n* and *K* given by random numbers, we performed simulation of the NARX model using the recovered data, ARX parameters obtained until step iii, and the given combination of *n* and *K*.

We calculated the Akaike information criterion (AIC) from the residual sum of squares between the experiment and simulation, number of parameters, and number of data to determine the parameters *n* and *K*. AIC is a measure of the relative quality of statistical models based on the trade-off between the goodness-of-fit of the model and the complexity of the model (Akaike, 1974). In step iv, we selected the combination of *n* and *K* with the minimum AIC and carried out signal recovery again using these *n* and *K*. We repeated steps i–iv 500 times, and selected the *n* and *K* and ARX parameters that minimize AIC for the training data set *AIC*_*training*_ in total (Figure 2B, step v). Let parameters with minimum *AIC*_*training*_ be parameters obtained from the training data set (Figure 2B, step v). Once these parameters were obtained, test data (still unequally spaced time series data) was added to the recovered Hankel-like matrix and signal recovery of the test data was performed (Figure 2B, step vi). With the parameters of the NARX model estimated from the training data set, we simulated the NARX model for test data LOO and calculated 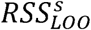, the residual sum of squares between experiment and simulation for test data set by stimulation condition *s* (NGF, PACAP, or PMA) (Figure 2B, step vii).

Because 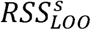 was obtained for each combination of training and test data set *s*, we took the sum of 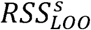 for test data set *s* as *RSS*_*LOO*_ (Figure 2B, step viii). We obtained the combination of *Inputs* as the identified *I*-*O* relationship that minimizes *RSS*_*LOO*_ for all combination of *Inputs* (Figure 2B, step ix). The *I*-*O* relationship indicates that a set of *Inputs* are selected as upstream molecules for each *Output*. In the final step, using this combination of input molecules, the parameters of the final NARX model were estimated by the procedure from step i to step v using all stimulation conditions as training data sets (Figure 2B, step x). Note that we used two different criterions *AIC*_*training*_ and *RSS*_*LOO*_; *AIC*_*training*_ to determine *n*, *k*, and ARX parameters, and *RSS*_*LOO*_ to select *Inputs* in order to save computational cost.

These estimated NARX parameters were used for further study (Figs. 3–6). The sensitivity with graded or switch-like response was obtained from the parameters of the Hill equation, and the gain and time constant were obtained from the parameters of the ARX model (Figure 2B, see “Calculation of Gain and Time Constant from the Linear ARX Model” section in the STAR METHODS). An example of the transformation of *Inputs* by the Hill equation and signal recovery following the ARX model and simulated *Output* is shown in Figure S2. We applied this method to identify the signaling-decoding system by gene expression underlying cell differentiation in PC12 cells using unequally spaced time series data with different time scales.

**Figure 3.**
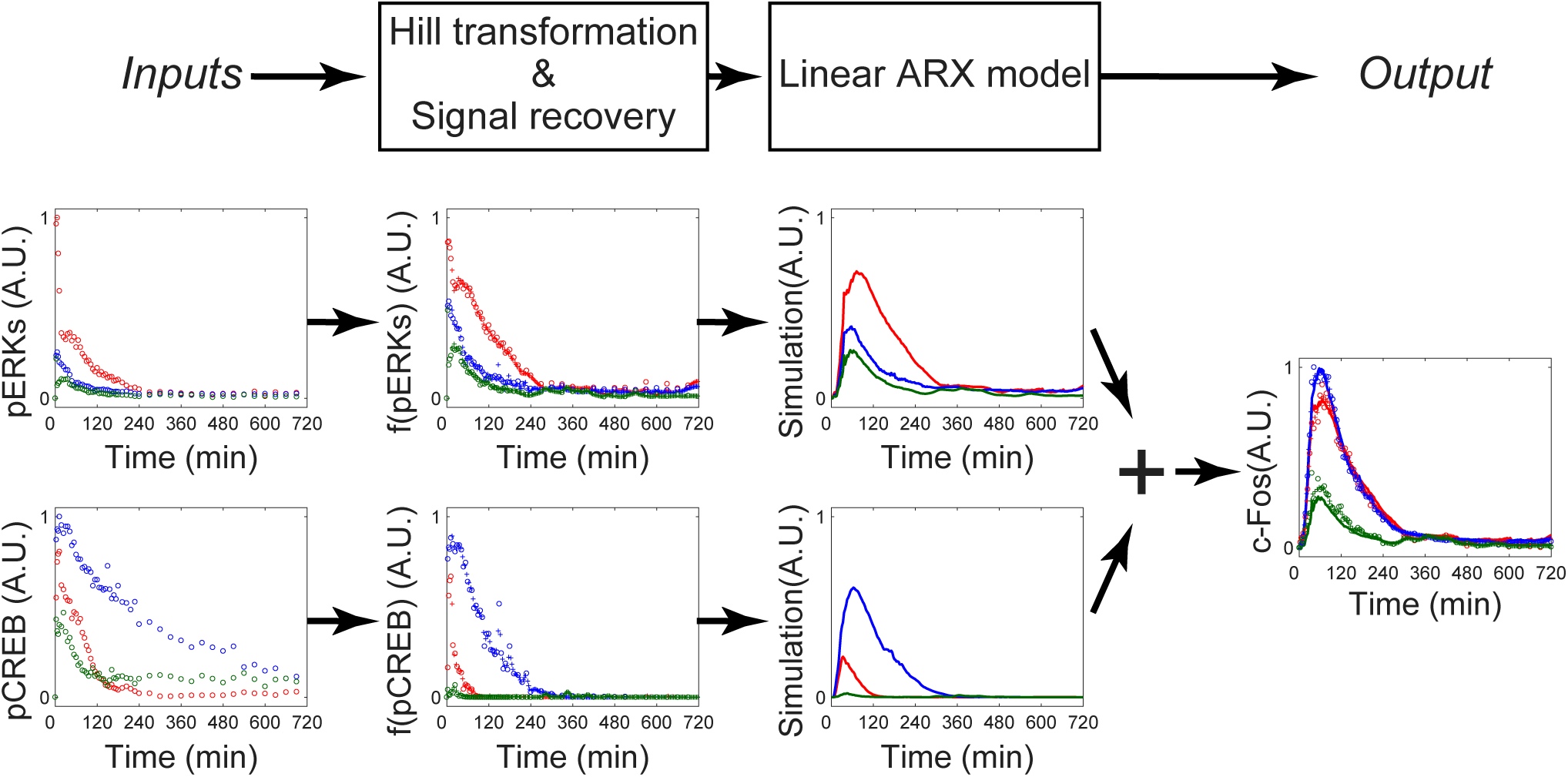
Transformation of *Inputs* by the Hill equation and signal recovery followed by the ARX model. Signal transformation in the nonlinear ARX model of c-Fos is shown. The signals of pERK and pCREB were transformed by the Hill equations and recovered. Then the transformed signals were temporally transformed by the linear ARX model. The sum of the transformed signals by the linear ARX model was c-Fos, the final output of the nonlinear ARX model of c-Fos.

### System Identification of Signaling-Dependent Gene Expression

We stimulated PC12 cells by NGF, PACAP, and PMA and measured the amount of phosphorylated ERK1 and ERK2 (pERK) and CREB (pCREB) and protein abundance of products of the IEGs, such as c-Jun, c-Fos, Egr1, FosB, and JunB by using QIC (Ozaki et al., 2010) (Figure 4). We chose these growth factors because they use different signaling pathways: NGF, PACAP, and PMA use Ras-, cAMP-, and PKC-dependent signaling pathways, respectively (Farah and Sossin, 2012; Gerdin and Eiden, 2007; Ravni et al., 2006; Vaudry et al., 2002). We also measured mRNA expression of LP genes such as *Metrnl*, *Dclk1*, and *Serpinb1a* using qRT-PCR (Figure 4). We measured the signaling molecules and gene expression with different sets of the time points because of the different time scales of temporal changes in signaling molecules and gene expression (Figure 4). Using the unequally spaced time series data with the different sets of the time points, we performed the system identification using integration of signal recovery and the NARX model (Figure 5A–C).

**Figure 5.**
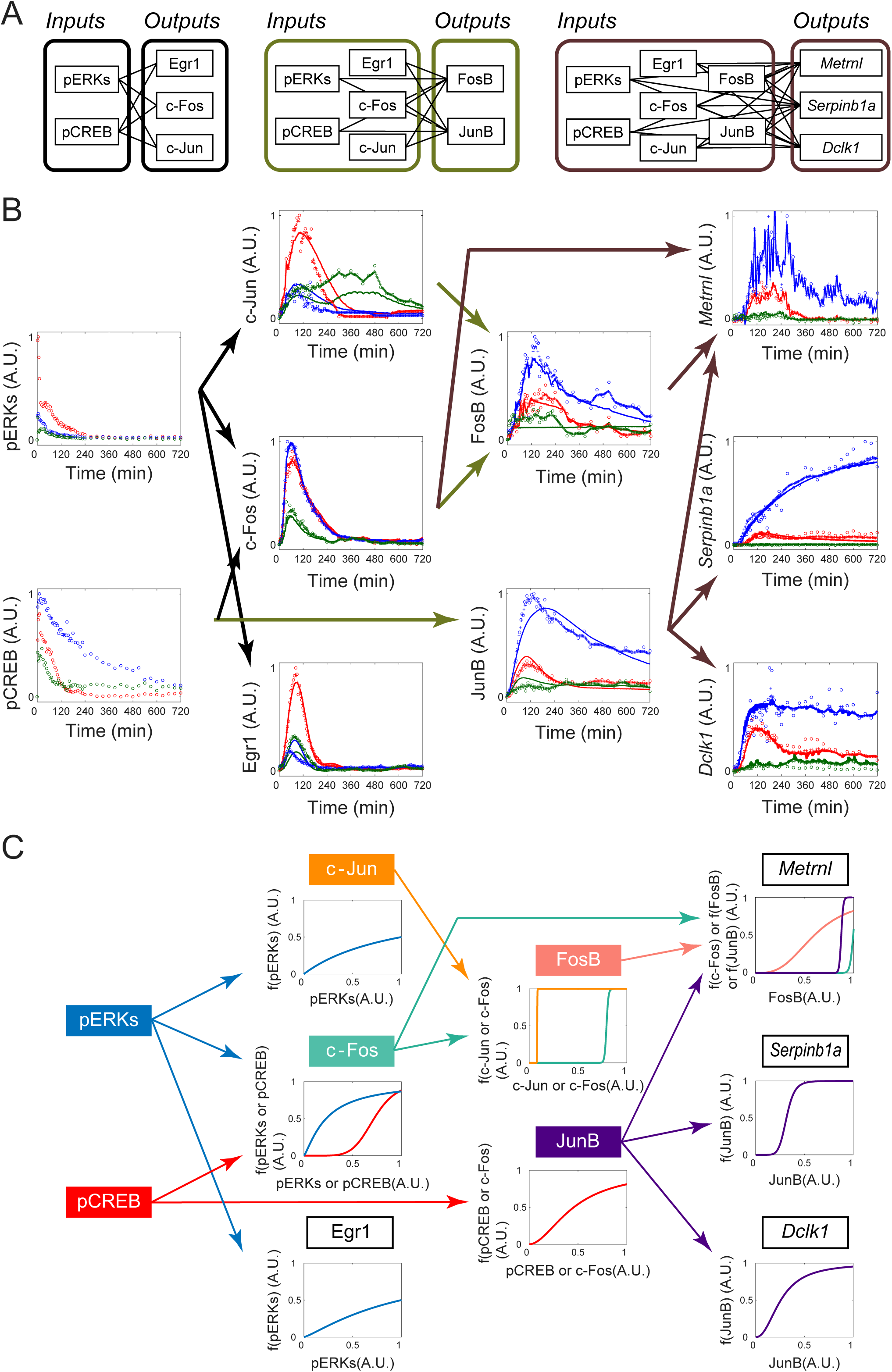
System identification of *I*-*O* relationships between the signaling, IEGs, and LP genes. (A) The sets of combinations of *Inputs* and *Outputs* for system identification. (B) The identified *I*-*O* relationships. Arrows indicate the estimated *I*-*O* relationships for each set of inputs and outputs. The colors of the arrows indicate the same sets of combinations of inputs and outputs as in (A). Dots, experimental data; pluses, the recovered signal data; solid lines, simulation data of the NARX model; red, NGF stimulation; blue, PACAP stimulation; green, PMA stimulation. (C) The dose-response curves obtained by the Hill equation. For each panel, conversion of *Inputs* by the identified Hill equation is shown. The colors of the arrows and plotted lines indicate the same *Inputs*, respectively.

**Figure 4.**
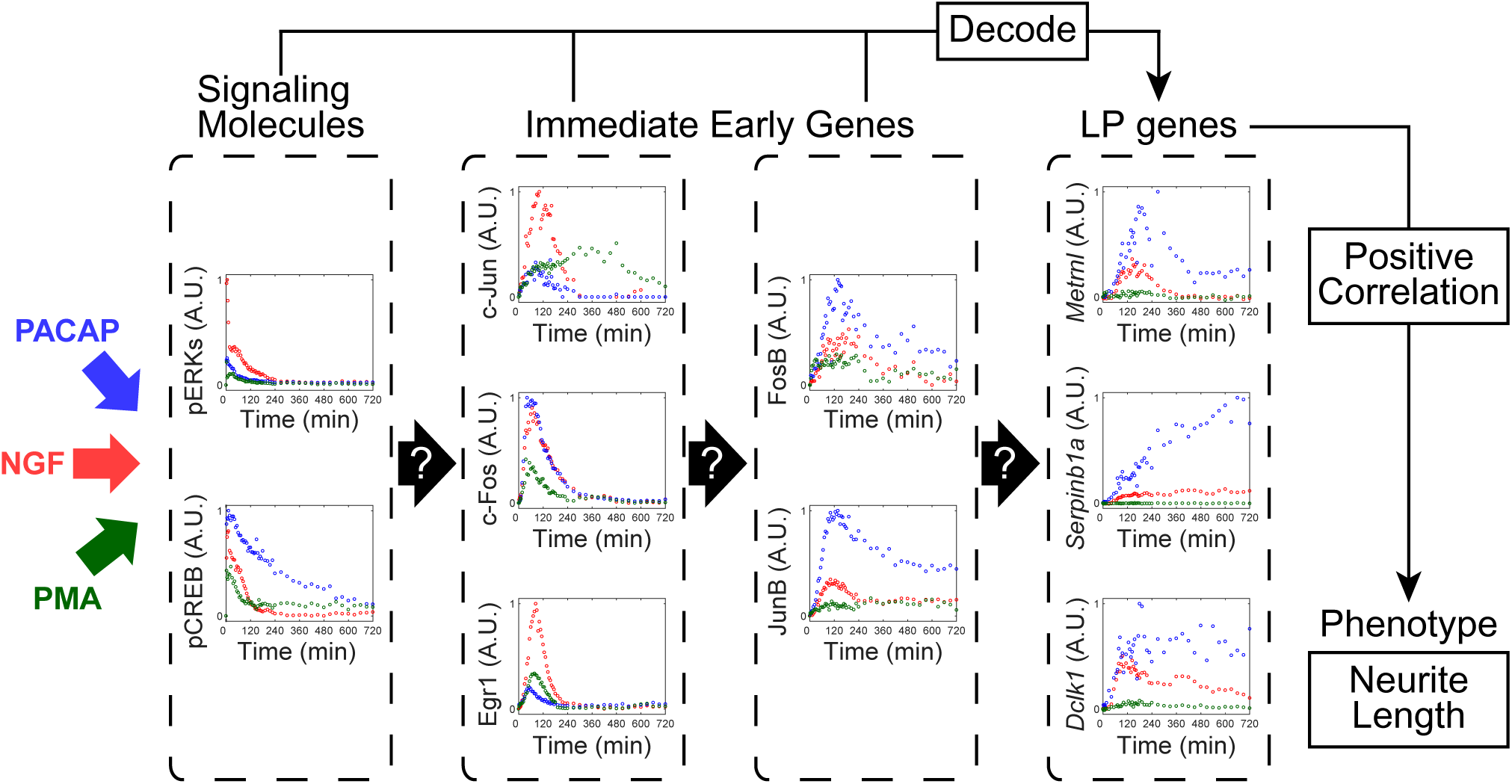
Experimental data of growth factor–dependent changes of signaling molecules and gene expression in PC12 cells. PC12 cells were stimulated with NGF (50 ng/ml, red), PACAP (100 nM, blue), or PMA (100 nM, green). Phosphorylation of signaling molecules, such as ERK and CREB, the product of IEGs such as c-Jun, c-Egr1, c-Jun, FosB, and JunB, and mRNA expression of LP genes such as *Metrnl*, *Dclk1*, and *Serpinb1a* were measured with different time points. These data were used for system identification by the NARX model in Figure 5–C.

Using these time series data sets, we selected three sets of *Inputs–Outputs* combinations from upstream to downstream and performed system identification for each set (Figure 5A). The system identification consists estimating the *I*-*O* relationship, dose-response by the Hill equation, and gain and time constant by the linear ARX model (Figure 3, see “Calculation of Gain and Time Constant from the Linear ARX Model” section in the STAR METHODS).

We selected pERK and pCREB as *Input* candidates for each *Output*, c-Jun, c-Fos, and Egr1, based on previous studies (Akimoto et al., 2013; Saito et al., 2013; Watanabe et al., 2012) (Figure 5A). We selected pERK, pCREB, c-Jun, c-Fos, and Egr1 as *Input* candidates for each *Output*, FosB and JunB (Akimoto et al., 2013; Saito et al., 2013; Watanabe et al., 2012) (Figure 5A). We selected pERK, pCREB, c-Jun, c-Fos, Egr1, FosB, and JunB as *Input* candidates for each *Output*, *Metrnl*, *Serpinb1a*, and *Dclk1* (Watanabe et al., 2012) (Figure 5A).

For c-Jun and Egr1, pERK was selected as an *Input*, and for c-Fos, pERK and pCREB were selected as *Inputs* (Figure 5). For FosB, c-Jun and c-Fos were selected as *Inputs*, and for JunB, pCREB was selected as *Input* (Figure 5B). For *Metrnl*, FosB, c-Fos and JunB were selected as an *Inputs*; however, contributions of c-Fos and JunB were negligible (Figure S2), indicating that FosB is a main *Input* for *Metrnl*. For *Serpinb1a* and *Dclk1*, JunB was selected as an *Input* (Figure 5B). It is noteworthy that FosB and JunB, but not signaling molecules and other IEGs, were mainly selected as *Inputs* of the LP genes and the inputs for *Metrnl* and *Dclk1* were different despite their similar temporal patterns.

We characterized the dose-response by the Hill equation and gain and time constant by the linear ARX model (Figure 5C, Table 1). The dose-responses from c-Jun and c-Fos to FosB showed typical switch-like responses, whereas others showed graded or weaker switch-like responses. Note that the gain from the converted c-Jun to FosB was much smaller than that from the converted c-Fos (Table 1), indicating that FosB is mainly regulated by c-Fos but not c-Jun.

**Table 1.**
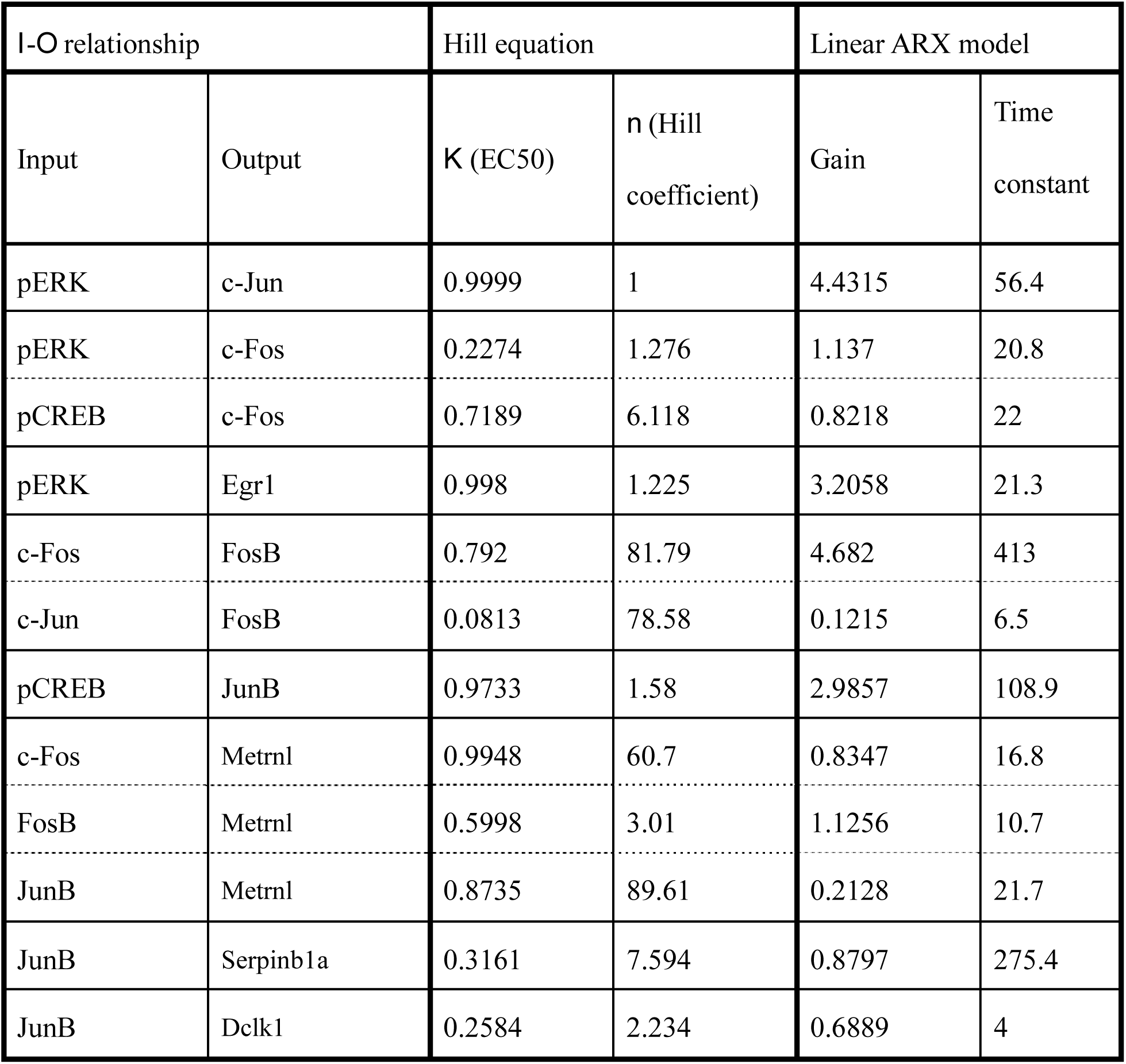
The identified *I*-*O* relationships, parameters of the Hill equation, and gain and time constant calculated from the linear ARX model in Figure 4.

The time constants for c-Jun, c-Fos, Egr1, *Metrnl*, and *Dclk1* were less than 1 h, whereas those for FosB, JunB, and *Serpinb1a* were more than 100 min (Table 1), indicating that induction of FosB and JunB temporally limit the overall induction of the LP genes from signaling molecules. The transformation of *Inputs* by the Hill equation followed by the ARX model is shown in Figure S2. In addition, when we integrated these three sets of the NARX model and simulated the response using only pERK and pCREB as *Inputs*, we obtained a similar result (Figure S3).

### Prediction and Validation of the Identified System by Pharmacological Perturbation

We validated the identified system by pharmacological perturbation. One of the key issues in PC12 cell differentiation is whether ERK or CREB phosphorylation mediates expression of the downstream genes (Ravni et al., 2006; Vaudry et al., 2002; Watanabe et al., 2012). Therefore, we selectively inhibited ERK phosphorylation by a specific MEK inhibitor, trametinib (Gilmartin et al., 2011; Watanabe et al., 2013; Yamaguchi et al., 2007) in PACAP-stimulated PC12 cells. We found that PACAP-induced ERK phosphorylation, but not CREB phosphorylation, was specifically inhibited by trametinib (Figure 6A, black dots).

**Figure 6.**
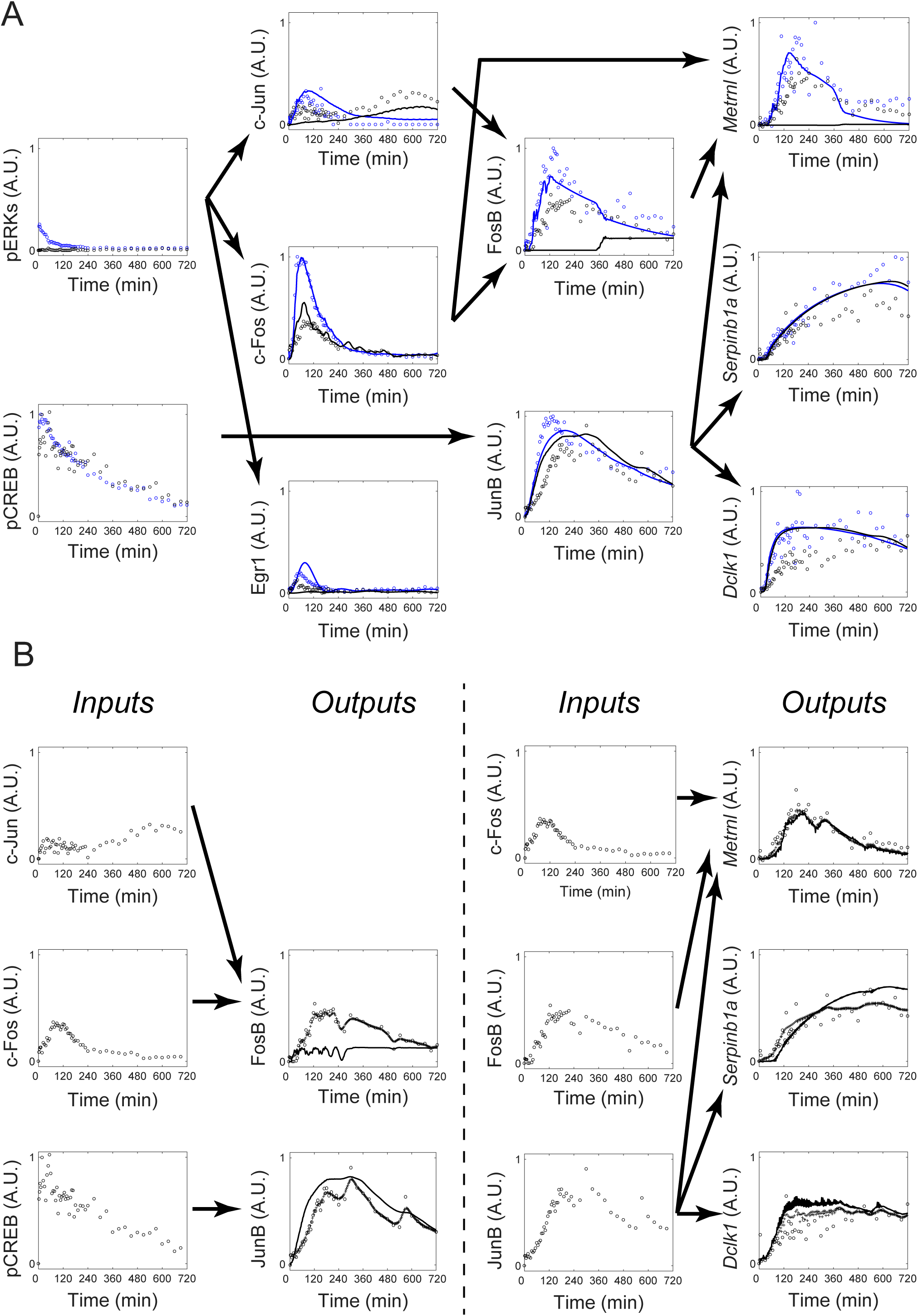
Prediction and validation of the identified system by pharmacological perturbation. (A) The predictive simulation result and experimental result by PACAP stimulation in the presence (black) or absence (blue) of trametinib. Lines, simulation; dots, experimental and recovered data. Experimental and recovered data of pERK and pCREB, and the simulated data of c-Jun, c-Fos, Egr1, FosB, and JunB are given as *Inputs*, and simulation was performed using the NARX model in Figure 5 (see Method). In the experiment, PC12 cells were treated in the absence (blue dots) or in the presence (black dots) of trametinib (10 μM) added at 30 min before stimulation with PACAP (100 nM). Note that the PACAP stimulation data are used, as in Figure 4. (B) Simulation using experimental and recovered data as *Inputs*. For each set of the *Inputs* (left panel for each) and *Outputs* (right panel for each), the unequally spaced time series data were recovered (pluses) (right panel for each), and the responses of *Outputs* were simulated by the NARX model identified in Figure 5A–C (solid lines) (right panel for each).

For c-Jun, c-Fos, Egr1, and JunB, we recovered signals of the unequally spaced time series data of *Inputs* and *Output*. For c-Jun, c-Fos, and Egr1, we simulated *Outputs* responses using these recovered data and the identified NARX model (Figure 6A black lines, see also “Simulation of the Integrated NARX Model” section in the STAR METHODS). For other downstream molecules, FosB, JunB, *Metrnl*, *Serpinb1a*, and *Dclk1*, we used the recovered data of pCREB and the simulated time series data of c-Jun, c-Fos, and Egr1 as *Inputs* for the identified NARX model (Figure 6A, black lines). The simulated time courses of *Outputs* were similar with those in experiments, except those of FosB and *Metrnl* (Figure 6A, black lines and black dots). In the simulation, FosB and *Metrnl* did not respond to PACAP in the presence of trametinib, whereas in the experiment both molecules did so, suggesting the possibility of failure of the system identification of FosB and/or *Metrnl*. Therefore, we investigated whether FosB and *Metrnl* can be reasonably reproduced when experimental and recovered data of c-Fos and c-Jun and of FosB, respectively, were used rather than the simulated ones (Figure 6B). When experimental and recovered data were used as *Inputs*, *Metrnl*, but not FosB, responded to PACAP in the presence of trametinib both in the simulation and experiment (Figure 6B), indicating that the failure of the system identification of FosB and *Metrnl* in Figure 6A arose from the failure of the system identification of FosB. Thus, all *Outputs* except FosB showed similar responses in the experiment and simulation when the experimental and recovered data were used as *Inputs*, indicating that in most cases the identified system is validated by pharmacological perturbation.

## DISCUSSION

In this study, we identified the system from signaling molecules to gene expression using the unequally spaced time series data for 720 min after the stimulation. Given that expression levels of the LP genes were highly correlated with neurite length regardless of growth factors (Watanabe et al., 2012) and expression continues for 720 min after the initial addition of NGF (Chung et al., 2010), the identified system is the selective growth factor–signaling decoding system for neurite length information, one of the most critical steps for cell differentiation in PC12 cells.

We previously identified the systems leading from pERK and pCREB to the IEGs using the equally spaced dense time series data with a uniform 3-min interval during 180 min (Saito et al., 2013). The identified *I*-*O* relationships in this study are the same, except for the inputs of FosB and JunB. In this study, for FosB c-Jun was selected as an *Input* in addition to c-Fos. However, the gain from the converted c-Jun to FosB was much smaller than that from the converted c-Fos (Table 1), indicating that the effect of c-Jun is negligible. For JunB, c-Fos was not selected as an *Input* in this study, whereas c-Fos was selected in the previous study (Saito et al., 2013). The gain from the converted c-Fos to JunB at lower frequency was much smaller than that from the converted pCREB (Saito et al., 2013), indicating that the effect of c-Fos is negligible in the previous study. Thus, the identified *I*-*O* relationships in this study are consistent with our previous work.

The estimated NARX parameters were also generally consistent with those in our previous study (Saito et al., 2013). The peak of c-Fos by NGF stimulation was approximately 0.9, whereas it was approximately 0.6 in our previous study (Saito et al., 2013). The difference may come from the difference in the algorithm for parameter estimation, because of the procedure of signal recovery is included in this study. Overall, the inferred the *I*-*O* relationship, the Hill equation, and the linear ARX model in this study are consistent with our earlier observations, indicating that system identification using unequally spaced time series data can give the same performance as using equally spaced time series data and that the system is time invariant during 720 min.

Furthermore, the identified system can reasonably reproduce the time series data using extrapolated data with trametinib, except for FosB. Taken together, these results demonstrate the validity of the predicted response of the identified systems. The reason for the failure of system identification of FosB is unclear, but it may reflect the failure of parameter estimation due to the insufficient number of experimental data points, the limitation of the NARX model structure, or the existence of unknown regulatory molecules. Further studies are necessary to address these issues.

One key issue is whether ERK and/or CREB mediates cell differentiation through downstream gene expression in PC12 cells (Ravni et al., 2006; Vaudry et al., 2002; Watanabe et al., 2012). We previously found that the LP genes are not induced by NGF in the presence of U0126, another MEK inhibitor (Watanabe et al., 2012), and that the MEK inhibitor blocks NGF-induced phosphorylation of both ERK and CREB in PC12 cells (Akimoto et al., 2013; Uda et al., 2013; Watanabe et al., 2012). By contrast, the MEK inhibitor blocked phosphorylation of ERK, but not CREB, in PACAP-stimulated PC12 cells (Akimoto et al., 2013; Uda et al., 2013) (Figure 6), suggesting that PACAP induces phosphorylation of CREB through a cAMP-dependent pathway, rather than the ERK pathway. These results demonstrate that NGF selectively uses the ERK pathway, whereas PACAP selectively uses the cAMP pathway for induction of the LP genes. Considering that LP genes are the common decoders for neurite length in PC12 cells regardless of growth factors (Watanabe et al., 2012), the identified system in this study (except for FosB) reveals the selective NGF- and PACAP-signaling decoding mechanisms for neurite length information.

Recently, fluorescence resonance energy transfer probes, optogenetics, and microfluidic devices have been developed to achieve observation and time control of ERK phosphorylation temporal patterns. These methods allow us to focus on quantitative relationships between various ERK phosphorylation temporal patterns and phenotypes such as cell differentiation (Albeck et al., 2013; Aoki et al., 2013; Doupé and Perrimon, 2014; Ryu et al., 2015; Sumit et al., 2017; Toettcher et al., 2013; Zhang et al., 2014). Although the relationship between signal transduction and phenotype has been extensively studied, it remains unclear how the signaling molecules quantitatively regulate the downstream gene expressions over a longer time scale, leading to cell fate decisions. In this study, we revealed the quantitative regulatory mechanism between signaling activation at a short time scale (tens of minutes) and gene expression at a longer time scale (day) by using a system identification method integrating a signal recovery technique and the NARX model based on compressed sensing.

A linear or spline interpolation is often used to convert unequally spaced time course data into equally spaced time course data in biological data analysis. However, such interpolation methods are not likely to be reliable because the interpolation methods ignore biochemical property of molecular network. By contrast, the interpolation used in this study is based on the NARX model, which reflects biochemical property. Thus, the proposal method in this study is biologically more plausible than a linear or spline interpolation.

There are obscure points for an application of this method to biological data analysis. The relationship between a number of observed time points and accuracy of signal recovery is theoretically unknown. In addition, how to select time points is also unknown. Intuitively, dense time points may be required for transient response, while sparse time points may be sufficient for sustained response. Further study is necessary to address this issue.

In molecular and cellular biology, molecular networks—the *I*-*O* relationship in this study—are generally examined by gene disruption or pharmacological perturbation experiments, meaning that the *I*-*O* relationship is examined using static and qualitative data. In this study, we used *Inputs*–*Outputs* time series data for system identification, allowing us to determine the *I*-*O* relationship using dynamic and quantitative data. Our system identification method does not require detailed knowledge of pathway information, which means that it can be used as a pathway finder directly from time series data. Moreover, additional information such as sensitivity with graded or switch-like response, time delay, and gain can be obtained. One of the advantages of using time series data rather than static perturbation experiments is simultaneously obtaining the *I*-*O* relationship, sensitivity, and time constant, which characterize the system behavior. This is based on the idea that input–output time series data implicitly include information on the *I*-*O* relationship. However, we must note that the *I*-*O* relationship obtained by using this method may be an apparent relationship inferred from time series data and is not necessarily the direct *I*-*O* relationship. Therefore, the obtained *I*-*O* relationship should be validated by gene disruption or pharmacological perturbation experiments, as shown in Figure 6.

In conclusions, we have devised a system identification method using unequally spaced sparse time series data by signal recovery. Because of technical and budget limitations in biological experiments, it is generally difficult to obtain sufficient numbers of equally spaced dense time series data of molecules with different time scales. Thus far, system identification based on time series data has been limited to phenomena with similar time scales. However, our system identification method can solve this time-scale problem and can be applied to any biological system with different time scales, such as the cell cycle, development, regeneration, and metabolism involving ion flux, metabolites, phosphorylation, and gene expression.

## Author Contributions

T.T., K. Konishi, and S.K. conceived the project and designed the experiments. T.T., N.M., and K. Kunida performed the experiments. T.T., M.F., K. Kunida, S.U., and K. Konishi derived the system identification algorithm by integrating signal recovery and the NARX model. T.T., M.F., and K. Konishi analyzed the data. T.T., M.F., N.M., H.K., K. Konishi, and S.K. contributed reagents/materials/analysis tools. T.T., M.F., K. Konishi, and S.K. wrote the manuscript.

### Acknowledgements

We thank Takamasa Kudo and Kosuke Umeda for helpful discussions and technical advice. We thank our laboratory members Atsushi Hatano, Kaoru Ohashi, Miharu Sato, and Risa Kuribayashi for their critical reading of this manuscript, helpful discussions, and technical assistance with the experiments. The computational analysis of this work was performed in part with support of the super computer system of the National Institute of Genetics (NIG), Research Organization of Information and Systems (ROIS). This work was supported by the Creation of Fundamental Technologies for Understanding and Control of Biosystem Dynamics, CREST, of the Japan Science and Technology Agency (JST). T.T. (Tsuchiya) receives funding from a Grant-in-Aid for Japan Society for the Promotion of Science (JSPS) Research Fellow (#14J12344). M.F. (Fujii) receives funding from a Grant-in-Aid for Challenging Exploratory Research (#16K12508). K. Kunida receives funding from a Grant-in-Aid for Young Scientists (B) (#16K19028). S.U. (Uda) receives funding from a Grant-in-Aid for Scientific Research on Innovative Areas (#16H01551). H.K. (Kubota) receives funding from a Grant-in-Aid for Scientific Research on Innovative Areas (#16H06577). K. Konishi receives funding from a Grant-in-Aid for Scientific Research (B) (#15KT0021), and (C) (#15K00246), and Grant-in-Aid for Scientific Research on Innovative Areas (#16H01554).

## STAR □METHODS

### KEY RESOURCES TABLE

STAR METHODS KEY RESOURCES TABLE
| REAGENT or RESOURCE | SOURCE | IDENTIFIER |
| --- | --- | --- |
| Antibodies |  |  |
| Mouse anti-phospho-ERK1/2 (Thr 202/Tyr 204) | Cell Signaling<br>Technology | Cat: 9106; RRID:<br>AB_331768 |
| Rabbit anti-phospho-CREB (Ser 133) | Cell Signaling<br>Technology | Cat: 9198; RRID:<br>AB_2561044 |
| Rabbit anti-c-Jun | Cell Signaling<br>Technology | Cat: 9165; RRID:<br>AB_2130165 |
| Rabbit anti-c-Fos | Cell Signaling<br>Technology | Cat: 2250; RRID:<br>AB_2247211 |
| Rabbit anti-Egr1 | Cell Signaling<br>Technology | Cat: 4154; RRID:<br>AB_2097035 |
| Rabbit anti-FosB | Cell Signaling<br>Technology | Cat: 2251; RRID:<br>AB_2106903 |
| Rabbit anti-JunB | Cell Signaling<br>Technology | Cat: 3753; RRID:<br>AB_2130002 |
| Alexa 488 goat anti-mouse IgG (H+L) | Invitrogen | Cat: A11029;<br>RRID: AB_138404 |
| Alexa 546 goat anti-rabbit IgG (H+L) | Invitrogen | Cat: A11035;<br>RRID:AB_143051 |
| Chemicals, Peptides, and Recombinant Proteins |  |  |
| DMEM | Sigma | Cat: D6046 |
| Fetal bovine serum | Sigma | Cat: 172012 |
| Horse serum | Gibco | Cat: 16050-122 |
| poly-L-lysine | Sigma | Cat: P4707 |
| NGF | R&D Systems | Cat: 1156-NG |
| PACAP | Sigma | Cat: A1439 |
| PMA | Sigma | Cat: P1585 |
| Trametinib | Selleckchem | Cat: S2673 |
| Can Get Signal immunostain solution A | Toyobo | Cat: NKB-501 |
| Hoechst 33342 | Invitrogen | Cat: H-3570 |
| Power SYBR Green PCR Master Mix | Applied Biosystems | Cat: 4367659 |
| Critical Commercial Assays |  |  |
| Agencourt RNAdvance Tissue Kit | Beckman Coulter | Cat: 32646 |
| High Capacity RNA-to-cDNA Kit | Applied Biosystems | Cat: 4387406 |
| Experimental Models: Cell Lines |  |  |
| PC12 rat adrenal pheochromocytoma cells | Masato Nakafuku (Ohio)<br>Sasagawa et al., 2005 | N/A |
| Oligonucleotides |  |  |
| Primers for qRT-PCR, see Table S2 | This paper | N/A |
| Software and algorithms |  |  |
| Iterative partial matrix shrinkage algorithm | Konishi et al., 2014 | http://dx.doi.org/10.1016/j.sigpro.2014.01.014 |
| MATLAB | MathWorks | https://www.mathworks.com/ |
| 7300 System SDS software version 1.3.1.21 | Applied Biosystems | N/A |
| Other |  |  |
| Biomek NX Span-8 liquid handling system | Beckman Coulter | N/A |
| Thermoshake heater-shaker | Variomag | Model number:<br>7100146-B |
| Robotic incubator STX-40 | Liconic | N/A |
| CellInsite NTX | Thermo Fisher Scientific | N/A |
| 7300 Real Time PCR System | Applied Biosystem | N/A |

### CONTACTS FOR REAGENT AND RESOURCE SHARING

Further information and requests for reagents and resources should be directed to and will be fulfilled by the Lead Contact, Shinya Kuroda.

### EXPERIMENTAL MODEL AND SUBJECT DETAILS

#### Cell Culture and Treatments

PC12 cells (kindly provided by Masato Nakafuku, Cincinnati Children’s Hospital Medical Center, Cincinnati, OH, USA) (Sasagawa et al., 2005) were cultured at 37°C under 5% CO_2_ in complete medium, Dulbecco’s modified Eagle’s medium (DMEM) (Sigma, Zwijndrecht, The Netherlands) supplemented with 10% fetal bovine serum (Sigma) and 5% horse serum (Gibco, Bethesda, MD, USA). For stimulation, PC12 cells were plated on poly-L-lysine-coated 96-well microplates (0.5×10^4^ cells/well) in the complete medium for 24 h and then treated with the complete medium in the presence or absence of the indicated doses of NGF (R&D Systems, Minneapolis, MN, USA), PACAP (Sigma), and PMA (Sigma) (Saito et al., 2013; Uda et al., 2013). Stimulations for cells seeded in 96-well microplates were performed by using a liquid handling system (Biomek NX Span-8, Beckman Coulter, Fullerton, CA, USA) with an integrated heater-shaker (Variomag, Daytona Beach, FL, USA) and robotic incubator (STX-40, Liconic, Mauren, Liechtenstein). For the inhibitor experiment, we stimulated cells with PACAP in the presence of 10 μM trametinib (Selleckchem, Houston, TX, USA). The inhibitor was added 30 min before stimulation with PACAP.

### METHOD DETAILS

#### Quantitative Image Cytometry

QIC was performed as previously described (Ozaki et al., 2010). Briefly, after stimulation by the growth factors, the cells were fixed, washed with phosphate-buffered saline, and permeabilized with blocking buffer (0.1% Triton X-100, 10% fetal bovine serum in phosphate-buffered saline). The cells were washed and then incubated for 2 h with primary antibodies diluted in Can Get Signal immunostain Solution A (Toyobo, Osaka, Japan). The cells were washed three times and then incubated for 1 h with second antibodies. After immunostaining, the cells were stained for the nucleus by incubating with Hoechst 33342 (Invitrogen, Carlsbad, CA, CA). The images of the stained cells were acquired by using a CellInsite NTX (Thermo Fisher Scientific) automated microscope with a 20× objective lens. For QIC analyses, we acquired different field images of the cells in each well, until the number of obtained cells exceeded 1000. Liquid handling for the 96-well microplates was performed using a Biomek NX Span-8 liquid handling system.

Intensities of the signaling activity and the IEGs between experiments were normalized by an internal control of each 96-well plate in QIC. Note that for the QIC assays, all the cells within a plate were fixed simultaneously to prevent the exposure of cells to formaldehyde vapor during the treatment.

#### qRT-PCR analysis

Reverse transcription–polymerase chain reaction (RT-PCR) was performed as previously described (Watanabe et al., 2012). Briefly, total RNA was prepared from PC12 cells using an Agencourt RNAdvance Tissue Kit according to the manufacturer’s instructions (Beckman Coulter, La Brea, CA, USA). RNA samples were reverse transcribed by using a High Capacity RNA-to-cDNA Kit (Applied Biosystems, Carlsbad, CA, USA) and the resulting cDNAs were used as templates for qRT-PCR. qRT-PCR was performed with Power SYBR Green PCR Master Mix (Applied Biosystems); the primers are shown in supplementary information Table S2. As an internal control for normalization, the β-actin transcript was similarly amplified using the primers. qRT-PCR was conducted using a 7300 Real Time PCR System (Applied Biosystems), and the data were acquired and analyzed by the 7300 System SDS software version 1.3.1.21 (Applied Biosystems). The sequences of the primers for the LP genes are shown in Table S2 (Watanabe et al., 2012).

#### NARX model and data representation

Assuming that the input molecules (*Input*) and output molecules (*Output*) signals satisfy the following NARX model, Eqs. (1) and (2), the system identification is performed by estimating unknown parameters in the NARX model,

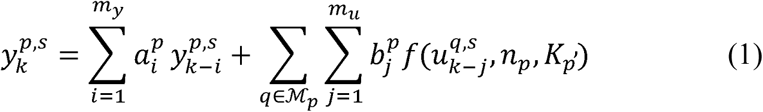

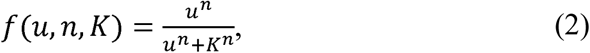

where 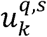 and 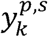 are experimental values of *Input* and *Output* at time step *k*, *p* and *q* respectively denote indices of *Output* and *Input* defined in the following sets,

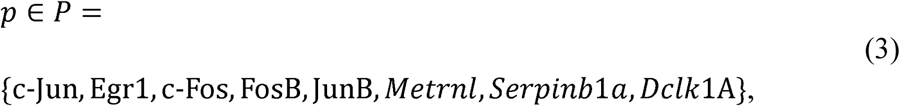

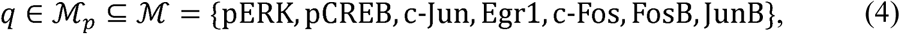

and *s* is an index of stimulation conditions defined as follows,

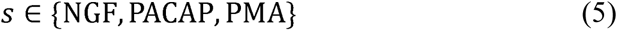

ℳ_*P*_ is the index set of *Input* defined for each *Output p ∈ P* (Fig. 5A). The nonlinear function *tcxJ* in Eq. (2) is the Hill equation that is one of the steady state solutions of biochemical reaction and widely used in the field of biology (Hill, 1910). The coefficients 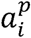 and 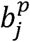, the orders *m*_*Y*_ and *m*_*u*_ in Eq. (1), *n*_*P*_ and *K*_*P*_ in Eq. (2), and set ℳ_*P*_ are unknown parameters. For each molecule under stimulation condition *s* (NGF, PACAP, and PMA), the unequally spaced time series data are obtained in this study. We consider them as equally spaced time data 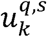 and 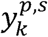 with missing time points and identify the unknown NARX parameters after recovering missing time points based on the low rankness of the Hankel-like matrix, which is described in the next section.

#### Extension of ARX System Identification from Unequally Spaced Time Series Data to the Nonlinear ARX System

To deal with the nonlinear ARX system, we extend ARX system identification from unequally spaced time series data to the nonlinear ARX system. First consider the simple case of the linear ARX model and then extend it to the NARX model. To perform system identification from unequally spaced time series data, equally spaced time series data are generated by signal recovery of unknown values of missing time points. Because the *Input* and *Output* data of a linear system are missing and the order of the system is unknown as in this study, the system identification using the recovered *Input* and *Output* data based on the low rankness of the Hankel-like matrix has been proposed (Liu et al., 2013). This is a method to recover missing data by solving the matrix rank minimization problem and to generate equally spaced sampled data.

In this study, we apply this matrix rank minimization approach to simultaneously identify the NARX model and recover missing data.

First, for simplicity, let us consider the case of the linear ARX model with single *Input* and single *Output* described by

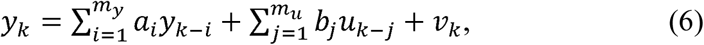

where *Y*_*k*_ and *u*_*k*_ are the *Output* and *Input* at time step *k*, and *V*_*k*_ is the noise. When only {*u*_*k*_}_*k*∈Ω_*u*__ and {*y*_*k*_}_*k*∈Ω_*y*__ are obtained, that is, the part of the *Input* and *Output* data 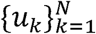 and 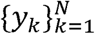, we consider the problem of recovering unknown *Input* and *Output* data. Here, *Ω*_*u*_ and *Ω*_*Y*_ are index sets and are a subset of the set {*1,2*,*…*, *N*}. We define Hankel-like matrices *Y* and *U* by Eqs. (7) and (8), where we assume that *N* is sufficiently larger than *r*.

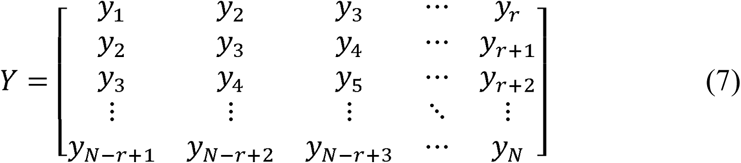

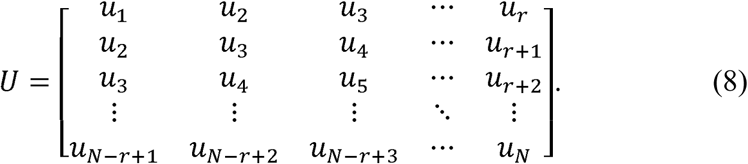

Hankel-like matrices *Y* and *U* are matrices called Hankel matrices if they are square matrices, and they are matrices in which the same components are entered from the lower left to the upper right in the matrix. Considering *V*_*k*_ *=0* in Eq. (6), that is, considering an ideal case without noise, Eq. (9) holds for the matrix [*Y U*] in which the matrices *Y* and *U* are arranged horizontally (Fig. S1B).

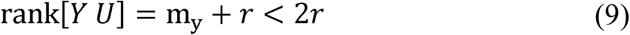

Thus, the matrix [*Y U*] is a low-rank matrix whose rank is determined by the order of the system. If *m*_*Y*_ is known in Eq. (9), the missing data can be recovered by restoring the unknown components of the matrix so that the rank of the matrix [*Y U*] becomes *m*_*y*_ *+ r*. Because the order *m*_*y*_ is unknown in this study, we recover the unknown components so as to minimize the rank of the matrix [*Y U*] based on the idea that it is better to describe the system with as few parameters as possible. That is, the missing data are recovered by solving the matrix rank minimization problem as follows,

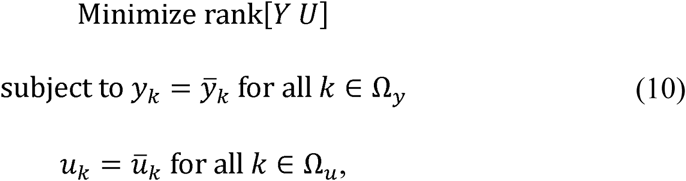

where 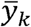 and 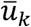 are observed values. Eq. (10) is a nonconvex optimization problem, which is generally a Non-deterministic Polynomial time (NP)-hard problem in the field of the computational complexity theory. Therefore, we relax this problem in Eq. (11) in which the objective function is replaced by the nucleus norm, the sum of the singular values of the matrix, and obtain a low-rank matrix by solving this optimization problem with the iterative partial matrix shrinkage (IPMS) algorithm (Konishi et al., 2014).

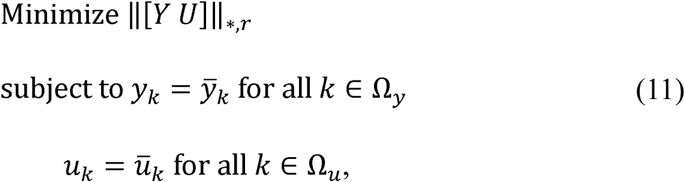

where ∥·∥_*,*r*_ represents the sum of singular values that are smaller than the *r*th greater singular value. The IPMS algorithm is a technique to provide a low-rank solution of Eq. (10) by solving Eq. (11) repeatedly for increasing *r* by 1, starting at *r=* 0, and provides recovered data with small energy loss after recovery and less distortion by preferentially estimating from a singular value of a large value (Konishi et al., 2014).

In the case of a multi-*Input* system, for each *Input*, a Hankel-like matrix *U*_*l*_ corresponding to the matrix *u* is generated, and by solving the matrix rank minimization problem of matrices arrayed side by side such as [*Y U*_1_… *U*_*L*_], *Inputs* and *Output* data can be similarly recovered. Also, when data under multiple stimulation conditions are obtained, *Input* and *Output* data can also be recovered by arranging the matrices vertically for each stimulation condition. For example, when there is a data set of NGF stimulation and PACAP stimulation and simulation condition *s* is *s ϵ* {*NGF*,*PACAP*}, a matrix composed of 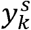 and 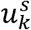 is vertically arranged for each stimulation condition *s* to construct *Y* and *U*, and *Input* and *Output* data can be recovered by solving the matrix rank minimization problem for [*Y U*].

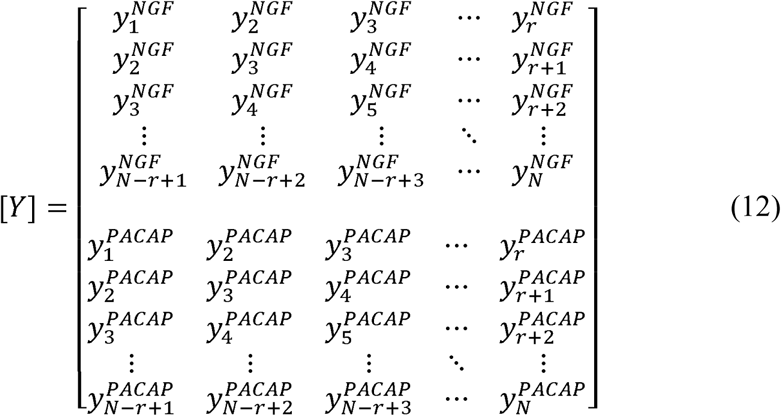

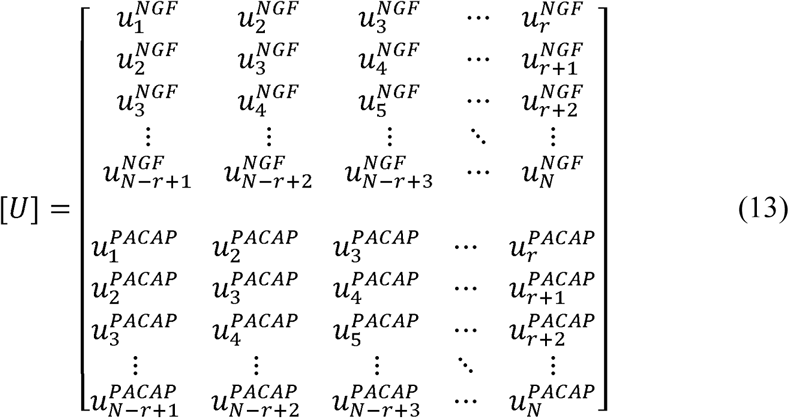

In the NARX model used in this study, because the observed *Input* data is nonlinearly transformed using the nonlinear function *f* in Eq. (2) and the *Input* data after transformation and the *Output* data follow the ARX system, signal recovery and system identification can be performed on the nonlinearly transformed *Input* data and *Output* data by the above method. Based on this idea, we performed nonlinear ARX system identification.

#### Procedure for System Identification by Integrating Signal Recovery and the NARX model

To estimate an *I*-*O* relationship, we prepare data sets of all combinations of input molecules (*Inputs*) for each output molecule (*Output*). For each data set, leave-one-out cross-validation is performed by preparing all combinations with only one test data set and the rest as the training data set. We have three stimulation conditions, NGF, PACAP, and PMA, and use two of them as the training data set and the other one as the test data set. Therefore, there are three combinations to divide the test and training data sets.

In nonlinear systems such as the NARX model in this study, even if all the *Input* and *Output* data are known, obtaining *n*_*P*_ and *K*_*P*_ is a nonconvex optimization problem, for which it is difficult to obtain an exact solution. Therefore, *n*_*P*_ and *K*_*P*_ are estimated by 500 trials with multiple random initial values. By repeating the following procedures from step i to step *v*, *n*_*P*_ and *K*_*P*_ are estimated so as to minimize the AIC for the training data set, while *Inputs* and *Output* of the NARX model are recovered. Subsequently, signal recovery of the test data set is performed in step vi, and the residual sum of square (RSS) is calculated for a test data set in step vii. Step vii is performed with all three combinations of training and test data sets, and take the sum of RSS for test data sets. Step viii is performed with all combinations of *Input*, and then in step ix a combination of *Input* with the minimum sum of RSS for test data sets is selected. This combination of *Inputs* is used for the *I*-*O* relationship. Using the combination of the *Input* molecules in step x and the data set of all stimulation conditions as the training data set, we estimate the parameters of the NARX model, which is used as the finally obtained NARX model (Figures 5 and 6).

#### Step i: Nonlinear transformation of *Input* data by the Hill equation

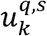, which is *Input q* at time step *k* under the stimulation condition *s*, is transformed into 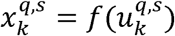 by Eq. (2), the Hill equation. The initial values of *n*_*p*_ and 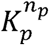 are given by *n*_*p*_ *=* 1 and a uniform random number between 0 to 1, respectively, for each *Input q*. Using the observed *Output 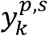* and the nonlinearly transformed *Input 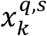,* the following Hankel-like matrix is constructed for *Output p* while assigning the previous closest observation value to the initial value of missing points. Note that this is a notation in the case of a single *Input*. Hereafter, two training data sets and one test data set are referred as *training*1 and *training* 2 and *test*, respectively.

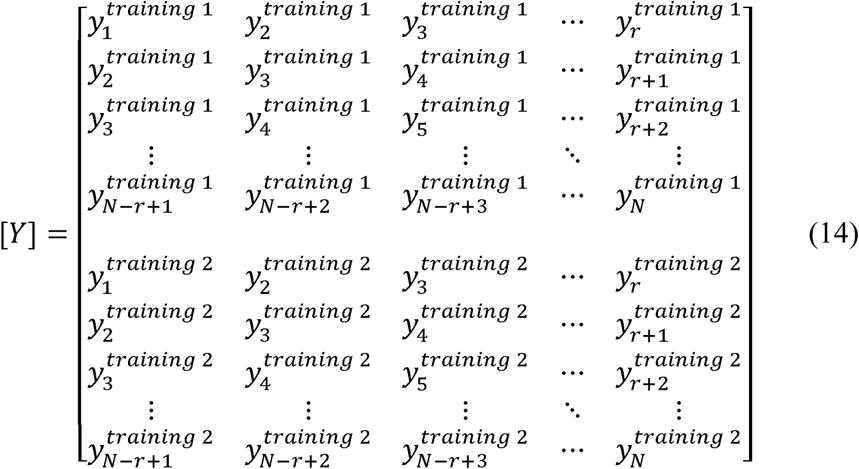

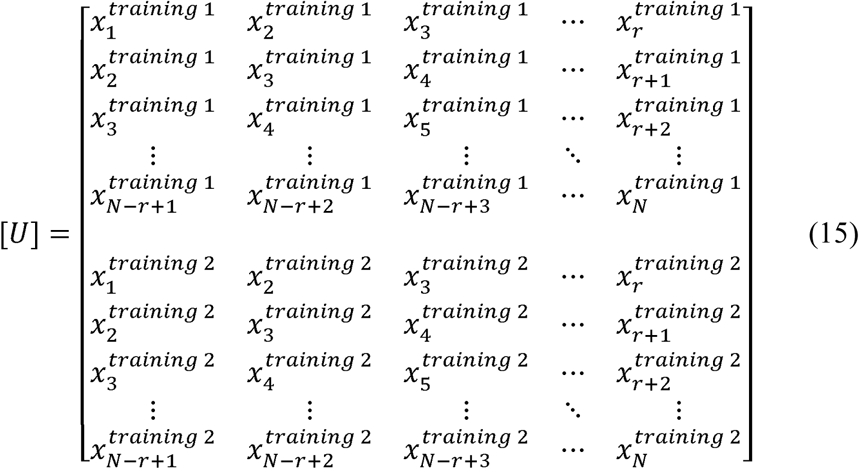

#### Step ii: Signal recovery of training data

Solve the matrix [*Y U*] rank minimization problem of Eq. (11) by the IPMS algorithm and recover converted *Input* data 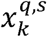 and *Output* data 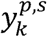. Note that, in the case of multi-*Input*, for each *Input*, a matrix *U*_*l*_ corresponding to the matrix *U* is generated, and by solving the matrix rank minimization problem of matrices arrayed side by side such as [*Y U*_1_… *U*_*L*_], *Inputs* and *Output* data can be similarly recovered.

#### Step iii: Calculate ARX parameters, *a* and *b*

Based on the relationship between the Hankel-like matrix and ARX parameters (Fig. S1B), obtain the ARX parameters 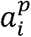 and 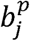 in Eq. (1) for *Output p* and each *Input q* using the recovered transformed *Input* data 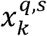 and *Output* data 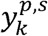. The order of the system, the lag order of the ARX model, is determined based on the matrix rank obtained in step ii.

#### Step iv: Estimate *n*_*p*_ and 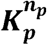 using the recovered data and ARX parameters

Using the inverse function *f* of Eq. (2), recover the missing time point data of *Input* before transformation by using Eq. (2). For the recovered 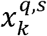 and 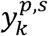, *n*_*P*_ in Eq. (2) is given again by uniform random numbers > 1 and ≤100 and 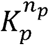 ≥0.001 and ≤1, and 200 combinations of *n*_*P*_ and 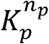 are generated. For each combination, perform simulation of the ARX model and calculate AIC for the training data set, *AIC*_*training*_. Select the combination of *n*_*P*_ and 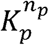 with the minimum *AIC*_*training*_. Using this *n*_*P*_ and *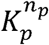*, *Input* and *Output* data in the matrix [*Y U*] composed of *Y* and *U* in Eqs. (14) and (15) is recovered again by the IPMS algorithm.

#### Step v: Select NARX parameters with the minimum *AIC*_*training*_

Repeat steps i to iv 500 times. Select *n*_*p*_ and 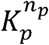 and ARX parameters that minimize *AIC*_*training*_.

#### Step vi: Signal recovery of test data

Using the *n*_*P*_, 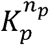 and ARX parameters selected in step v, add test data to the recovered matrix [*Y U*] in Eqs. (14) and (15) like in Eqs. (16) and (17). Test data are also recovered by solving the test data added matrix [*Y U*] rank minimization problem with the IPMS algorithm. Note that training data sets have already been recovered until step v, and signal recovery of only test data is performed in this step.

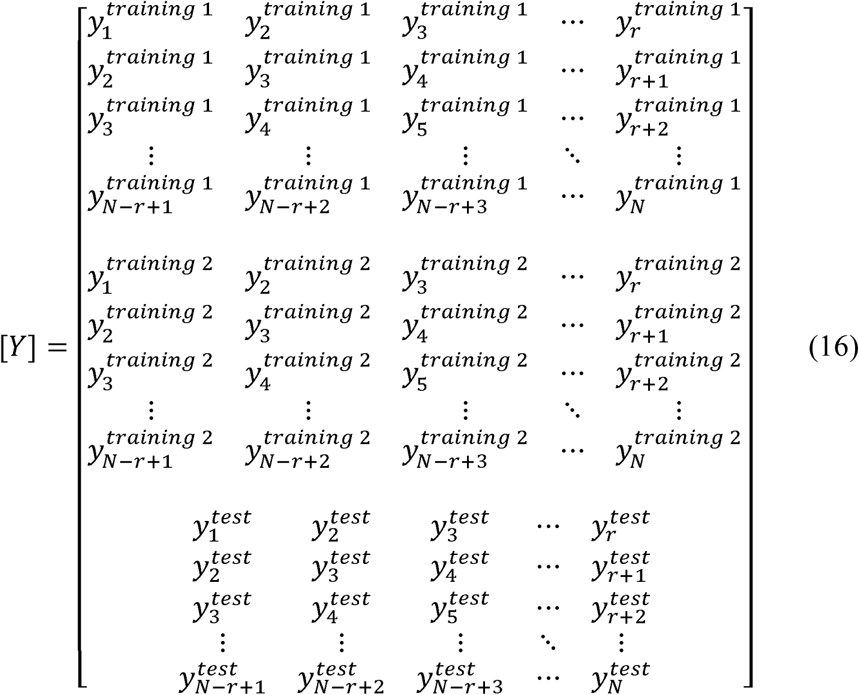

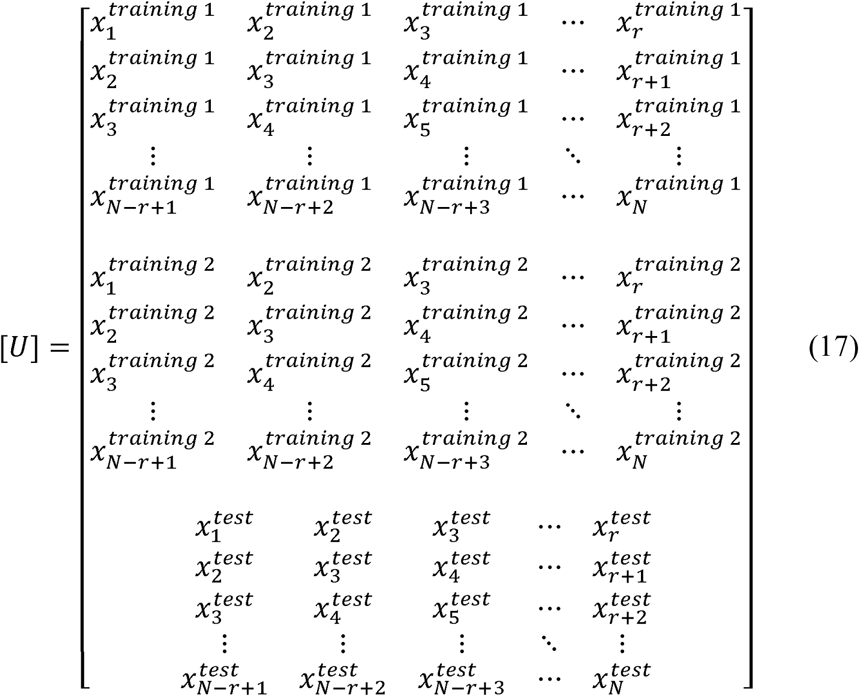

#### Step vii: NARX model simulation and calculate 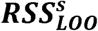 for test data set *s*

Simulate the test data using equally spaced time series data recovered in step vi and parameters of the Hill equation and ARX parameters. Calculate the 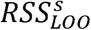, the residual sum of square for the stimulation condition of the test data set.

#### Step viii: Calculate *RSS*_*LOO*_ by taking the sum of 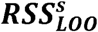 for each stimulation *s*

Perform steps i to vii for all three combinations of training and test data sets. Let *RSS*_*LOO*_ be the sum of 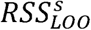 for each stimulation *s* of test data set.

#### Step ix: Obtain the *Input* combination with minimum *RSS*_*LOO*_

Perform steps i to viii for all combinations of *Inputs*. Select the combination of *Inputs* with the minimum *RSS*_*LOO*_ for the *I-O* relationship.

#### Step x: Estimate the NARX model with signal recovery using all data sets

Using the combination of *Input* determined in step ix, estimate the NARX parameter with signal recovery by the procedure from steps i to v using all stimulation conditions as training data sets.

Note that, when simulating with the ARX model, set the value of *Output* to 0 before time 0, otherwise the value of the *Output* obtained by the simulation is used to obtain the next time value. For stimulation in Figure 6, signal recovery was performed by step vi using experimental data with trametinib as the test data set.

#### Calculation of Gain and Time Constant from the Linear ARX Model

Gain and time constant *τ* were calculated from the frequency response function obtained from the linear ARX model. For simplicity, we consider here the case of a single *Input* single *Output* ARX model like Eq. (6), which can be re-described as follows,

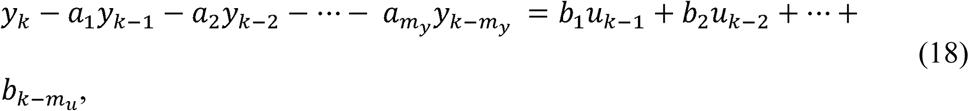

and its *Z*-transform are given by

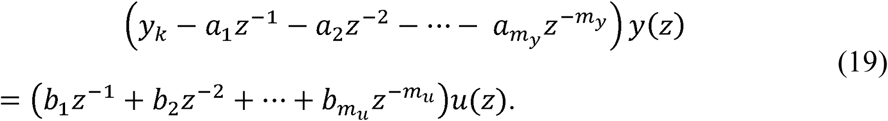

Then a discrete-time transfer function, a function to convert *Input* to *Output* through the system, *G*(*z*) can be described using these ARX parameters,

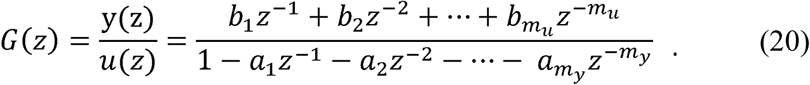

To consider the frequency response function and calculation of gain and phase, *z* is substituted by *iω*,

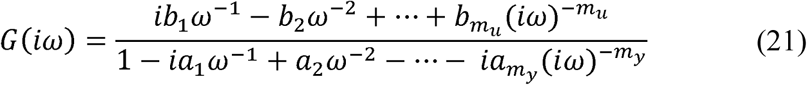

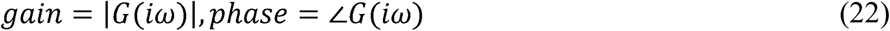

where *i* is an imaginary unit and *w* is frequency. Therefore, gain and phase can be calculated from ARX parameters. The frequency response curve and phase diagram at each *Input* and *Output* of the identified linear ARX model are shown in Figure S4. Note that gain indicated in Table 1 is steady-state gain. From the frequency response function, cutoff frequency *f*_*cutoff*_, an inverse of time constant *τ*, is obtained by calculating the frequency at which the gain corresponds to 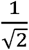 of the steady-state gain. Because Eq. (23) is established between *f*_*cutoff*_ and the time constant *τ*, we can obtain *τ* from the ARX parameters through the above procedure.

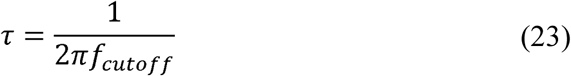

#### Simulation of the Integrated NARX Model

The simulation of the integrated NARX model was performed as follows. Experimental and recovered data of pERK and pCREB, and the simulated data of c-Jun, c-Fos, Egr1, FosB, and JunB were given as *Input* data and simulation was performed using the NARX model in Figure 5.

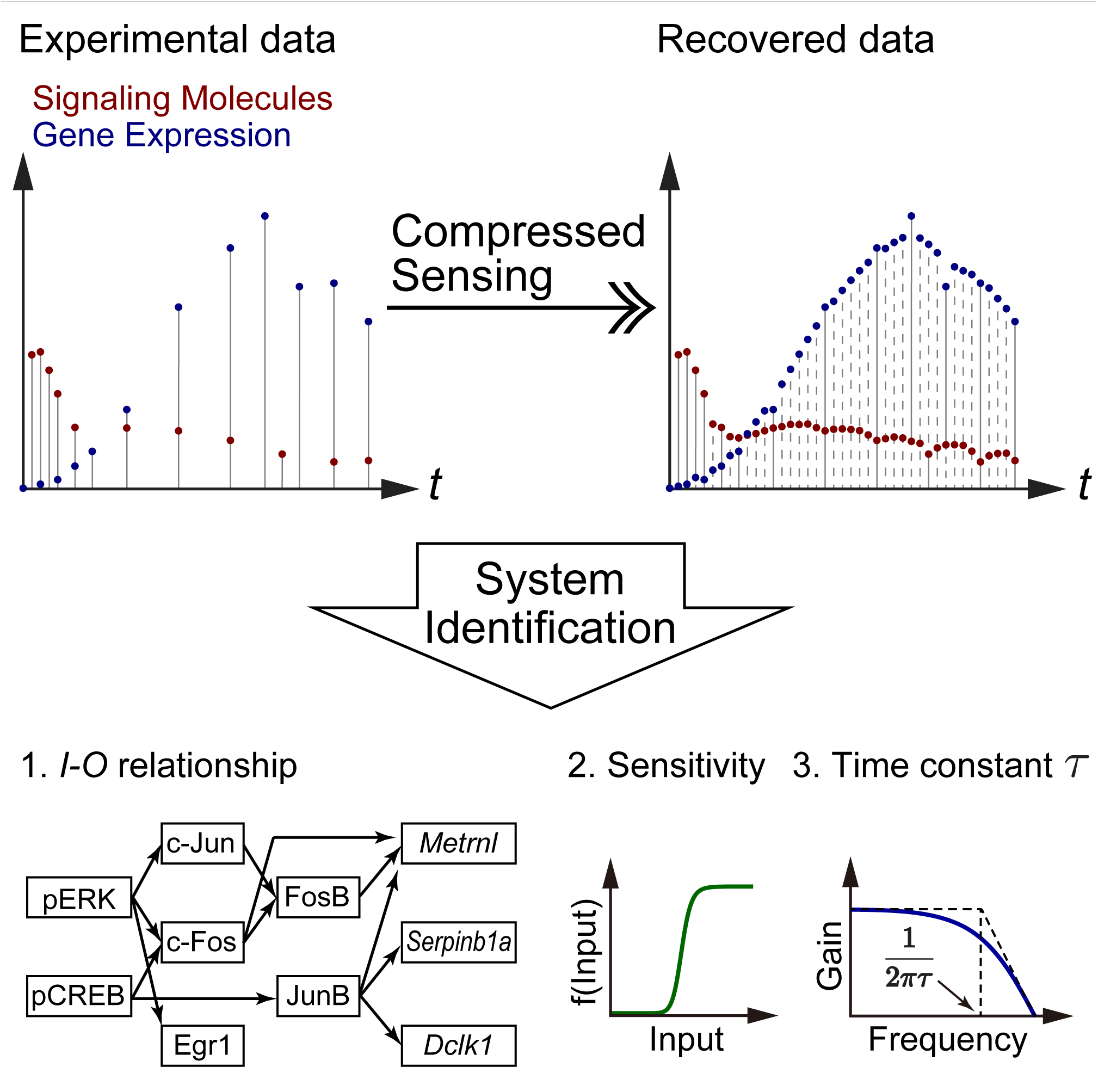

